# Simultaneous Morphology, Motility and Fragmentation Analysis of Live Individual Sperm Cells for Male Fertility Evaluation

**DOI:** 10.1101/2021.05.11.443542

**Authors:** Keren Ben-Yehuda, Simcha K. Mirsky, Mattan Levi, Itay Barnea, Inbal Meshulach, Sapir Kontente, Daniel Benvaish, Rachel Cur-Cycowicz, Yoav N. Nygate, Natan T. Shaked

## Abstract

We present a new technique for sperm analysis on individual unstained live cells, measuring DNA fragmentation, morphology with virtual staining, and motility. The method relies on quantitative stain-free interferometric imaging and deep-learning frameworks, and is utilized for male fertility evaluation. In the common clinical practice, only motility evaluation is carried out on live human cells, while full morphological evaluation and DNA fragmentation assays require different staining protocols, and therefore cannot be performed on the same cell, resulting in inconsistencies in fertility evaluation. We use a clinic-ready interferometric module to acquire dynamic sperm cells without chemical staining, and deep learning to evaluate all three scores per cell. We show that the number of cells that pass each criterion separately does not accurately predict how many would pass all criteria. This stain-free evaluation method is expected to decrease uncertainty in infertility diagnosis, increasing treatment success rates.

## 1. Introduction

Infertility is a common condition, as roughly one in six couples face some degree of infertility, defined as failing to naturally conceive over the course of one year. Male infertility factors contribute to approximately half of all cases.^[1]^ In vitro fertilization (IVF) has made parenthood possible for many people that could not conceive otherwise. IVF enables fertilization of a female egg by a male sperm outside the female body. The selection of sperm cells is crucial especially for intracytoplasmic sperm injection (ICSI), a common type of IVF, in which the clinician chooses a single sperm cell using a micropipette and injects it into the female egg in a dish. Approximately 10 eggs are obtained in each IVF cycle through a surgery subsequent to the woman’s hormonal treatment, and it is not uncommon that no fertilized eggs develop into good-quality embryos.^[2]^ Unfortunately, much less efforts are invested in analyzing sperm cells than eggs for IVF sperm selection. The World Health Organization (WHO) provides criteria for male fertility evaluation based on imaging the sperm sample.^[3]^ These criteria mainly consist of evaluation of the sperm morphology, motility, and DNA fragmentation status. While motility assessment is done on live sperm cells swimming in a dish, internal morphological assessment and DNA fragmentation assays are typically performed on fixed and stained sperm cells, and on different portions of the sample; hence, different evaluations are done on different cells. Certain sperm cells may possess good morphology, but not good motility, or may possess both good morphology and motility, but fragmented DNA, causing defective embryo development or delivery.^[4]^ Current tests cannot distinguish between these scenarios leading to a lack of consistency in sperm evaluation and selection performed by different clinicians, as well as a large margin of error, even when automated analysis is performed.

Deep convolutional neural networks (CNNs) have recently proven to be an efficient tool for image analysis and classification.^[5-9]^ The model is generally built from sequential layers, each providing a nonlinear mapping of the previous layer output to the following layer. Recently, generative adversarial neural networks (GANs)^[10]^ have been successfully used for virtual staining of microscopic images.^[11-15]^ These neural networks include a generator model and a discriminator model, where the generator takes random noise and maps it to an image, and the discriminator classifies images as real or generated.^[16]^ Deep learning automatic classification of sperm cells could be the next gold standard. However, it is still difficult to obtain reliable automatic sperm classifiers or virtual staining using only qualitative two-dimensional images as an input. The biological mechanisms that connect sperm movement, morphology, and contents to fertilization potential and normal pregnancy is not fully understood yet.^[17,18]^

In the absence of staining, sperm cells are nearly transparent under bright-field microscopy, as their optical properties differ only slightly from those of their surroundings, resulting in a weak image contrast. An internal contrast mechanism that can be used when imaging sperm cells without staining is their refractive index.^[19-21]^ Phase imaging creates stain-free quantitative image contrast based on the optical path delay (OPD) induced in the light beam as it interacts with the sample, which can be recorded by interferometry. Conventional phase contrast imaging methods for sperm cells, such as Zernike phase contrast microscopy, differential interference contrast (DIC) microscopy, and Hoffman modulation contrast microscopy are not quantitative, thus they do not enable interpretation of the resulting phase images in terms of the quantitative optical thickness of the sperm cell. In addition, these techniques present significant imaging artifacts, especially near the cell edges. Quantitative phase imaging records the full sample complex wave front including the optical thickness map of the cell, which is equal to the integral of the refractive-index values across the cell thickness. This map is proportional to the cell dry mass surface density, thereby providing cellular parameters that have not been available to clinicians.^[22]^ Until recently, quantitative phase imaging implementations were limited to optics labs, due to the optical setup bulkiness, difficulty of alignment, and sensitivity to mechanical vibrations. In recent years, we made significant efforts and succeeded to make these wave front sensors applicable and affordable for direct clinical use.^[21,23-25]^

DNA fragmentation is a critical biomarker in sperm cells. Studies have shown that even in normal semen samples, approximately 20% of the sperm cells have fragmented DNA, which becomes worse with age.^[26]^ Sperm DNA fragmentation has been associated with reduced fertilization rates, reduced embryo quality, reduced pregnancy rates and increased miscarriage rates. Thus sperm cells with fragmented DNA should not be selected for IVF. DNA fragmentation is not well correlated with morphology and motility^[27]^ and cannot be imaged at the moment without staining. Detecting DNA fragmentation requires molecular staining, which cannot be carried out during IVF.

In this work, we present a new approach for measuring DNA fragmentation status of live and unstained dynamic sperm cells, at the individual cell level, in parallel to measuring the cell motility and morphology as though they have been stained. This is achieved by using a clinic-ready interferometric module to acquire dynamic sperm cells and record their quantitative phase profiles, followed by using algorithmic architecture that includes CNNs to virtually stain the cells and classify them based on their morphology, motility, and DNA fragmentation status. All information is then mapped to a 3-D scatter plot for each cell, thereby characterizing male fertility better than has been possible until now. In contrast to previous attempts of finding correlation between pairs of stain-free bright-field (low-contrast) images and DNA-fragmentation-stained images using machine learning,^[28,29]^ here we use rapid interferometric imaging, providing not only the amplitude image, but also the quantitative phase profile of the cell. Since the phase profile is quantitative, having meaningful content-related values on all the cell points, the gap between the images in the input pair is smaller; thus, the presented approach allows better individual-cell DNA fragmentation classification, in addition to internal-morphological-structure virtual staining of each cell (which is not possible by bright-field imaging), enabling better fertility grading, based on the triple generalized score (WHO morphology, motility and DNA fragmentation). Totally, we analyze 51,809 human sperm images and show that the number of cells that pass all three criteria cannot be accurately determined by the number of cells that pass each criterion separately, which necessitates a different fertility grading procedure than used today.

## 2. Results

The architecture of the overall system for attaining a generalized fertility evaluation per patient is shown in **Figure 1**. The proposed technique enables simultaneously obtaining, for each of the dynamic sperm cells imaged in a dish, the full stain-like morphology, DNA fragmentation, and motility status, without the need for chemical staining. We implemented a clinic-ready holographic setup (**Figure S1**) to acquire stain-free quantitative phase maps of the cells. This setup is comprised of an inverted microscope and a custom-built common-path, compact interferometric module, connected to the microscope camera port (see Methods). The acquired off-axis interferograms were processed into the optical thickness maps of the sperm cells, taking into consideration both their physical thickness and their refractive index contents. As shown in **Figure S2** and in **Video S1**, these maps can be visualized in HEMA-staining-like colors, thereby distinguishing between different sub-cellular structures. We designed a custom-built tracking algorithm and tracked all cells dynamically, resulting in a space-time array per cell. We then obtained the full intracellular morphology and motility parameters, as well as the DNA fragmentation status, and mapped each cell to one point in a 3-D space, with the axes being morphology, motility, and fragmentation values, thereby displaying the complete cellular status of each cell. The combined cell points per patient in the 3-D space are depicted as a sphere, with the sphere volume representing the patient’s generalized fertility score.

**Figure 1.**
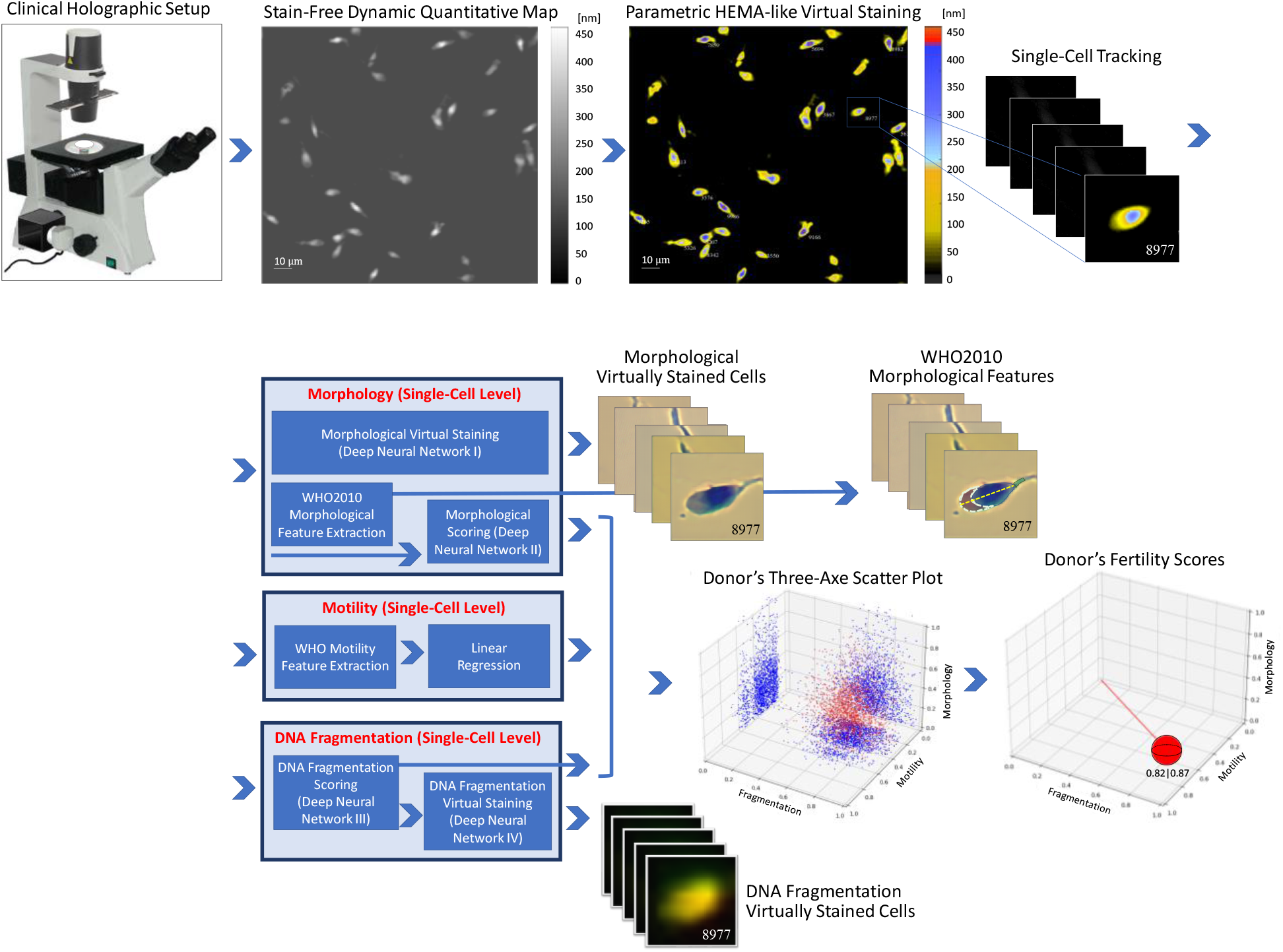
System schematics and working modes. Architecture for analyzing the full sperm morphology, motility, and DNA fragmentation status of large numbers of dynamic live sperm cells at the single-cell level, using stain-free quantitative imaging and deep learning, yielding more accurate donor fertility scores.

Using this scheme, we quantitatively imaged 51,809 human sperm images of 5,101 cells from eight donors. The collection of all cells measured per donor and their representation on the 3-D scatter plot uniquely characterizes the fertility status of the donor. Instead of having three different criteria measured on three different populations of the sample, where there is the risk of having cells that pass one criterion and fail one or more of the other criteria, we are now able to generate a triple-criteria score per cell.

### 2.1. Full Morphology Evaluation and Virtual Staining

As shown in **Figure 1** and **Video S1**, the morphological evaluation of each cell includes both virtual staining of the cell, extraction of the cell morphological features, and the final cell classification, as recommended by the 2010 World Health Organization^[3]^ (WHO2010) for stained cells, although no chemical staining is used and the cells are not fixated but rather dynamically swim in a dish. For both the cell virtual staining and the WHO2010-based morphology classification tasks, we used deep neural networks. For virtual morphological staining, we used the model presented by us.^[14]^ In short, we trained a deep neural network with the stain-free optical thickness maps of sperm cells and their chemically stained counterparts. Then, the trained network could transform th\e stain-free optical thickness maps into virtually stained images, making them look as though they were chemically stained, without actual staining (**Figure S3**), thereby providing the information necessary for gold-standard evaluation. The network was tested by classification of a trained embryologist, after randomizing the data order, and yielded very similar results to those obtained by chemical staining. In the current study, virtual staining is implemented on live and dynamic sperm cells for the first time. We further analyzed the cells using classical image processing techniques to automatically extract six morphological features including nucleus area, acrosome area, total head area, mean posterior-anterior difference, dry mass, and the variance of the optical thickness values quantifying the texture of the cell.^[30]^ We then trained another deep neural network to classify sperm cells based on their WHO2010 morphology criteria. During the training process, the network receives the optical thickness map of the cell along with the ground-truth WHO2010 classification of a trained embryologist, and outputs six values; the five WHO2010 criteria per cell (head shape, acrosome size, number of vacuoles, midpiece shape and orientation, and cytoplasmic droplets) together with a direct overall prediction of whether the cell passes all five criteria (independent of the first five outputs) according to the embryologist. In addition, a combined overall prediction checking if the model predicted a ‘pass’ for all five qualifications presented by the first five outputs was calculated, for comparison with the direct overall prediction (see Methods). **Figure 2** presents the architecture of the neural network used, as well as the Receiver operating characteristic curve (ROC) and precision recall curve (PRC). The ROCs for the five qualifications attained areas under the curves (AUCs) of 95.7%, 96.3%, 100.0%, 98.7% and 86.7%, respectively. The PRCs for the five qualifications attained AUCs of 93.6%, 95.5%, 100.0%, 99.7% and 97.4%, respectively. The direct overall parameter, representing the direct classification, attained AUC of 96.2% for the ROC and 93.5% for the PRC, precision of 90.9%, and accuracy of 93.1%. Precision is a dominant predictor for a successful sperm classifier, as a person’s potential of fertilization is mostly dependent on his best cells. These cells reach the egg in natural insemination, and are the only cells that should be selected for IVF. Once the network is trained and tested, it can classify the cell without having the ground truth label from the trained embryologist. The direct overall parameter was set as the coordinate value of the morphological axis in the 3-D scatter plot for the cell examined.

**Figure 2.**
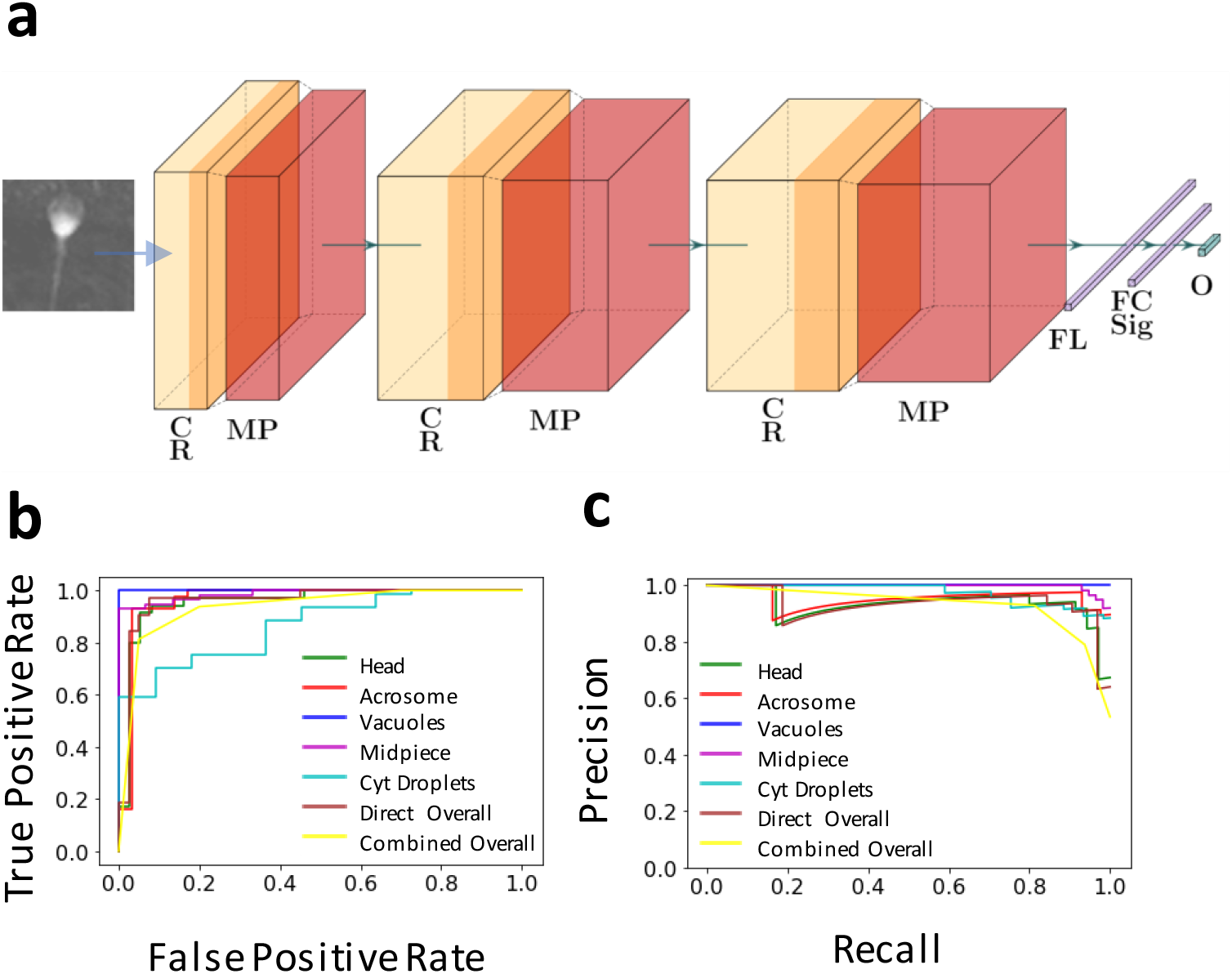
Deep neural network for morphological sperm classification. a) CNN architecture predicting morphological features per stain-free image. b) ROC curves for the seven outputs of the network. The direct overall score is a prediction whether the cell passes all five criteria (independent of the first five outputs), and the combined overall score indicates whether the first five output are positive. c) Precision-recall curves for the seven outputs of the network. C – Convolutional layer, R – ReLU activation, MP – Max pooling layer, FL – Flattening layer, FC – Fully connected layer, Sig – Sigmoid activation function, O – Output.

### 2.2 Motility Evaluation

To assess the sperm motility on an individual-cell basis, we used the space-time arrays extracted from the cell tracking algorithm, comprised of the locations of each cell in each frame. The trajectory for one representative cell is shown in **Figure 3**. We divided the trajectory into one-second windows with a half window stride, as the qualification of swimming linearly in progressively motile cells is only expected over short time intervals. One of these windows is shown on the right of **Figure 3**. We then calculated eight motility parameters suggested by the WHO2010. These include: curvilinear velocity (VCL), straight-line velocity (VSL), average path velocity (VAP), linearity (LIN), wobble (WOB), straightness (STR), beat-cross frequency (BCF), and mean angular displacement (MAD). The median values of these parameters over all windows per cell were taken as the final motility parameters per cell. For each donor, another sub-sample was tested by an experienced embryologist according to the qualitative WHO2010 protocol, classifying each sperm cell into one of three motility classes: immotile, non-progressively motile and progressively motile. To compare the qualitative and quantitative motility tests, we conducted two comparative experiments. First, we calculated the correlation between the qualitative and quantitative motility test results over all eight donors. To do this, we chose four quantitative motility parameters (VCL, VSL, VAP and VSL×LIN), which most resembled the qualitative assessment, resulting in high significant correlations. To cancel the effect of sampling errors, a video containing 87 sperm cells was processed both quantitatively by our automatic algorithm and qualitatively by the embryologist. This increased the correlations (from 0.49 to 0.75) and their statistical significance (p-values decrease from 0.15 to 10^−16^). We then used a least-square approximation to define a linear equation that maps all automatically extracted quantitative motility parameters to the three qualitative classes defined above, to ensure that no previously acquired fertility score is overlooked by our protocol. The normalized function value of the linear equation was used as the coordinate of the motility axis in the 3-D scatter plot.

**Figure 3.**
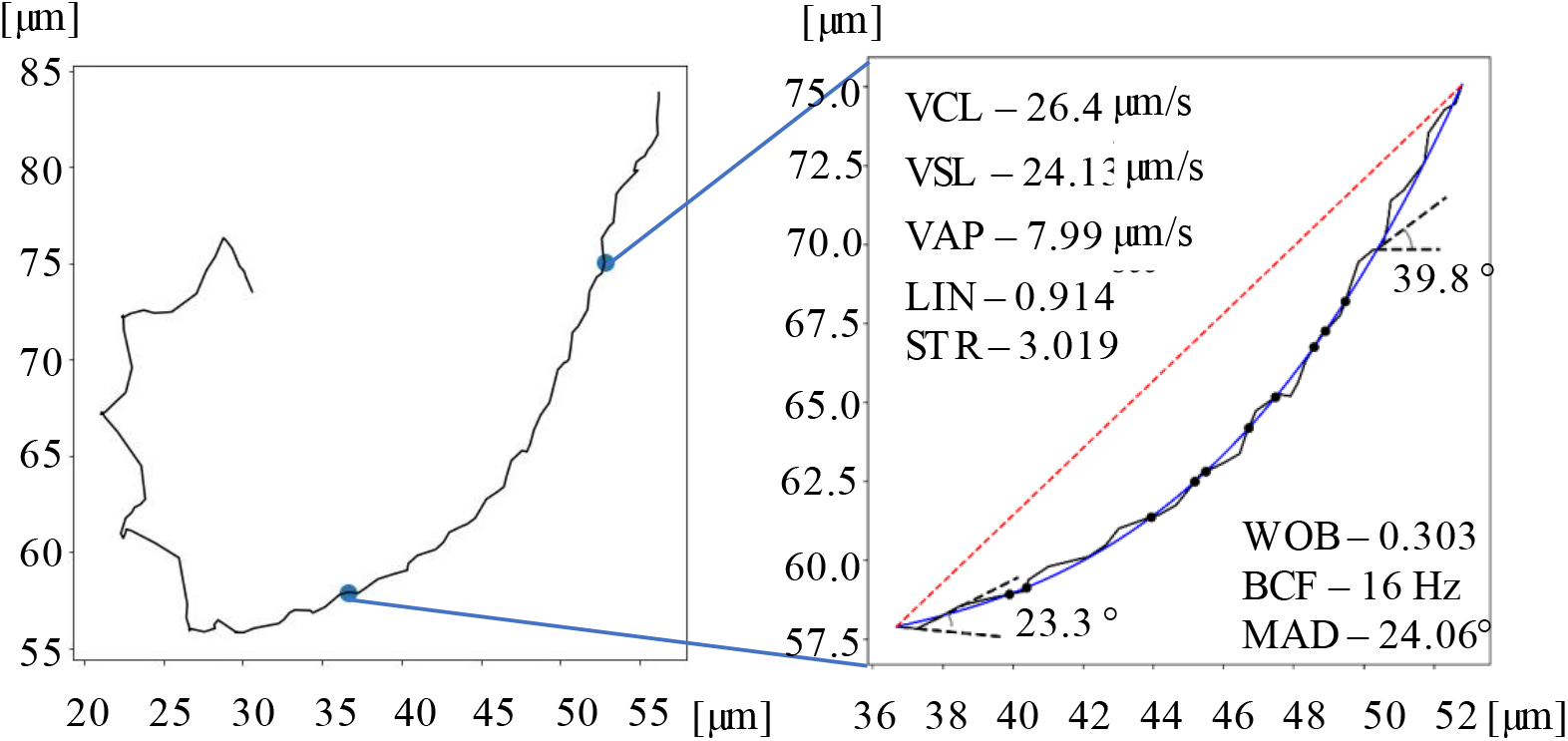
Automatic motility feature extraction. Example of a sperm cell tracked over 4.9 sec (left), enlargement of a 1-sec segment and automatic calculation motility parameters (right).

### 2.3 DNA Fragmentation Evaluation and Virtual Staining

We used a deep neural network to grade each live sperm cell according to its DNA fragmentation level. In contrast to Barnea et al.^[31]^ that shows only statistical difference for specific parameters in large populations of sperm cells measured by interferometry, with population overlaps, here we provide an individual-cell DNA measurement without staining. For training, we used pairs of images: the stain-free quantitative thickness map of the cell and the image of the same cell after it was stained by acridine orange, a DNA fragmentation indicator emitting green fluorescence for double-stranded DNA (‘non fragmented’) and red fluorescence for single-stranded DNA or RNA (‘fragmented’). Results are shown in **Figure S4**. The automatically extracted morphological parameters were also inserted into the network, creating a bimodal neural network. **Figure 4a** presents the network architecture (see also Methods). The accuracy, precision, ROC AUC and PRC AUC were all above 0.98. After training and testing, the network can take a stain-free quantitative optical thickness map of the cell and output the cell fragmentation level without the need for chemical staining. Another deep neural network was trained to virtually stain the quantitative optical thickness map, creating a semblance of its acridine-orange-stained counterpart. The network architecture is shown in **Figure 4b** and its virtual staining operation is demonstrated in **Figure 4c**, yielding a virtually stained image that is very similar to the chemically stained sperm cell image shown in the center.

**Figure 4.**
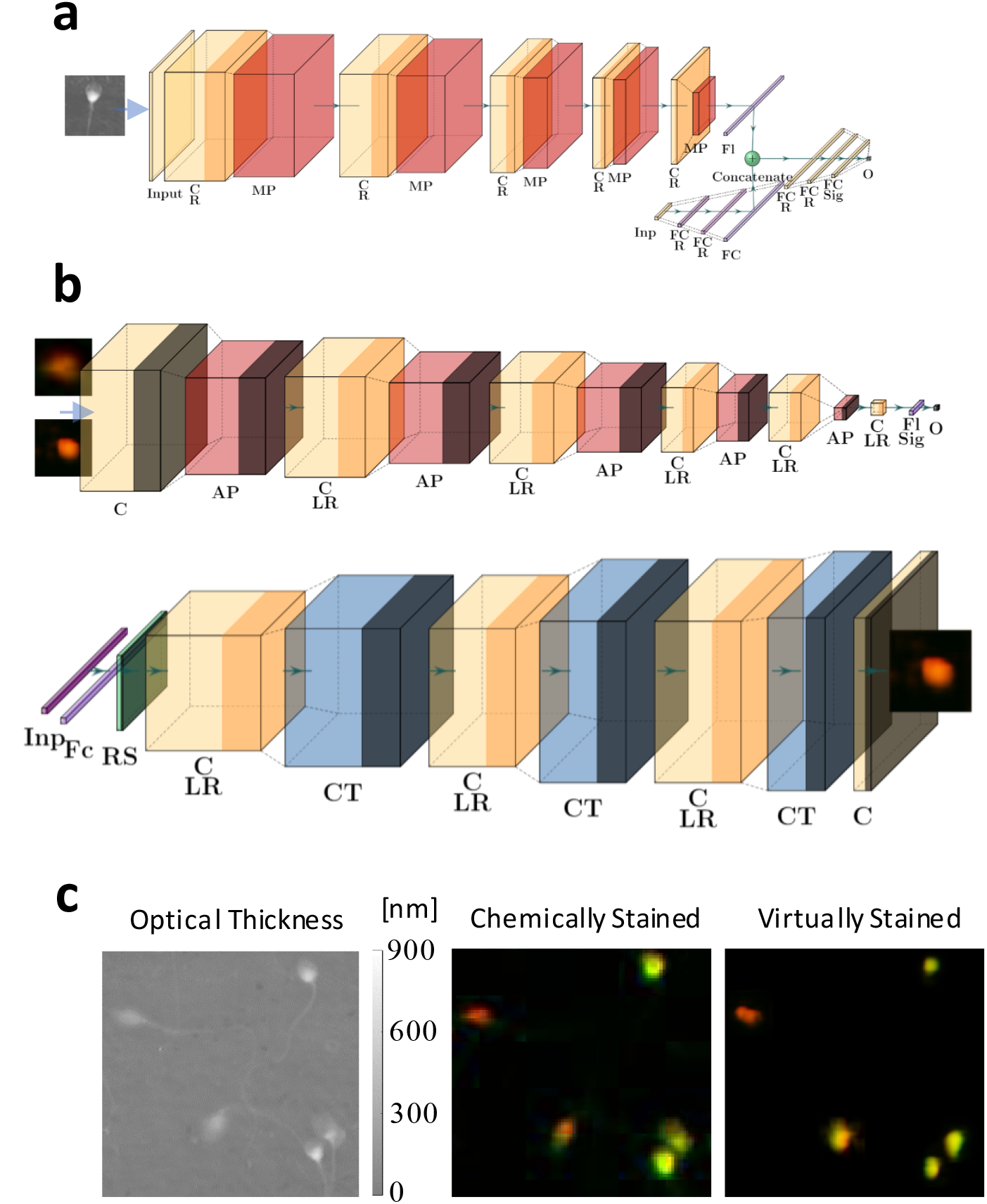
Stain-free DNA fragmentation classification and virtual staining using deep neural networks, working at the single cell level. a) Architecture of the bi-modal neural network, predicting the fragmentation level for each stain-free optical thickness map. b) Architecture for the GAN model used for DNA fragmentation virtual staining with the discriminator network (top) and generator network (bottom). c) Comparison between unstained optical thickness map (left) with the actual chemically stained cell image (center) and the generated virtually stained cell image (right). C – Convolutional layer, R – ReLU activation, MP – Max pooling layer, Fl – Flattening layer, FC – fully connected layer, Sig – Sigmoid activation function, O – Output, AP – Average pooling layer, LR – Leaky ReLU activation, RS – Resampling layer, CT – Convolution transpose layer.

### 2.4 Patient’s Fertility Scoring

To evaluate the patient’s fertility, we calculated each sperm-cell morphology, motility and DNA fragmentation as explained above, for live cells and without staining, and then mapped the cell into a single point in a 3-D space with axes representing these three criteria. Our evaluation also consists of virtual morphological and acridine-orange staining per cell. As shown in **Figure 5**, the presented technique enables gathering all single-cell triple-criteria information regarding the population of sperm cells. The left column in **Figure 5** shows the resulting 3-D scatter plots for the eight donors measured, while presenting both the 2-D projection of each cell on each of the three planes, and also the location of that cell in the 3-D space. The right column in **Figure 5** shows the intersections of the three criteria and the left column shows the Venn diagrams, where each of the three circles represents a specific criterion. The numbers of cells that pass each criterion, together with the number of cells that pass each set of two criteria and all criteria is displayed, where the passing thresholds are set to 0.5 for morphology, 0.29 for motility, and 0.61 for fragmentation. This figure demonstrates the great variability between donors. In relation to the other criteria, some donors have more cells that pass the fragmentation tests (Donors 2–5), while others have more cells that pass the motility test (Donors 1,6–8). Some donors have more cells that are both non-fragmented and morphologically intact (Donors 1–3,5,7), while others have more cells that are both morphologically intact and motile (Donors 4,6,8). The percentage of the overall passing cells differs among the donors: Donor5 (**Figure 5e**) has significantly more cells than Donor 2 **(Figure 5b**), yet Donor 2 has three more sperm cells that pass all three criteria. The thresholds may be adjusted according to the patient’s specific needs, as shown in **Figure S5** and **Video S2**.

**Figure 5.**
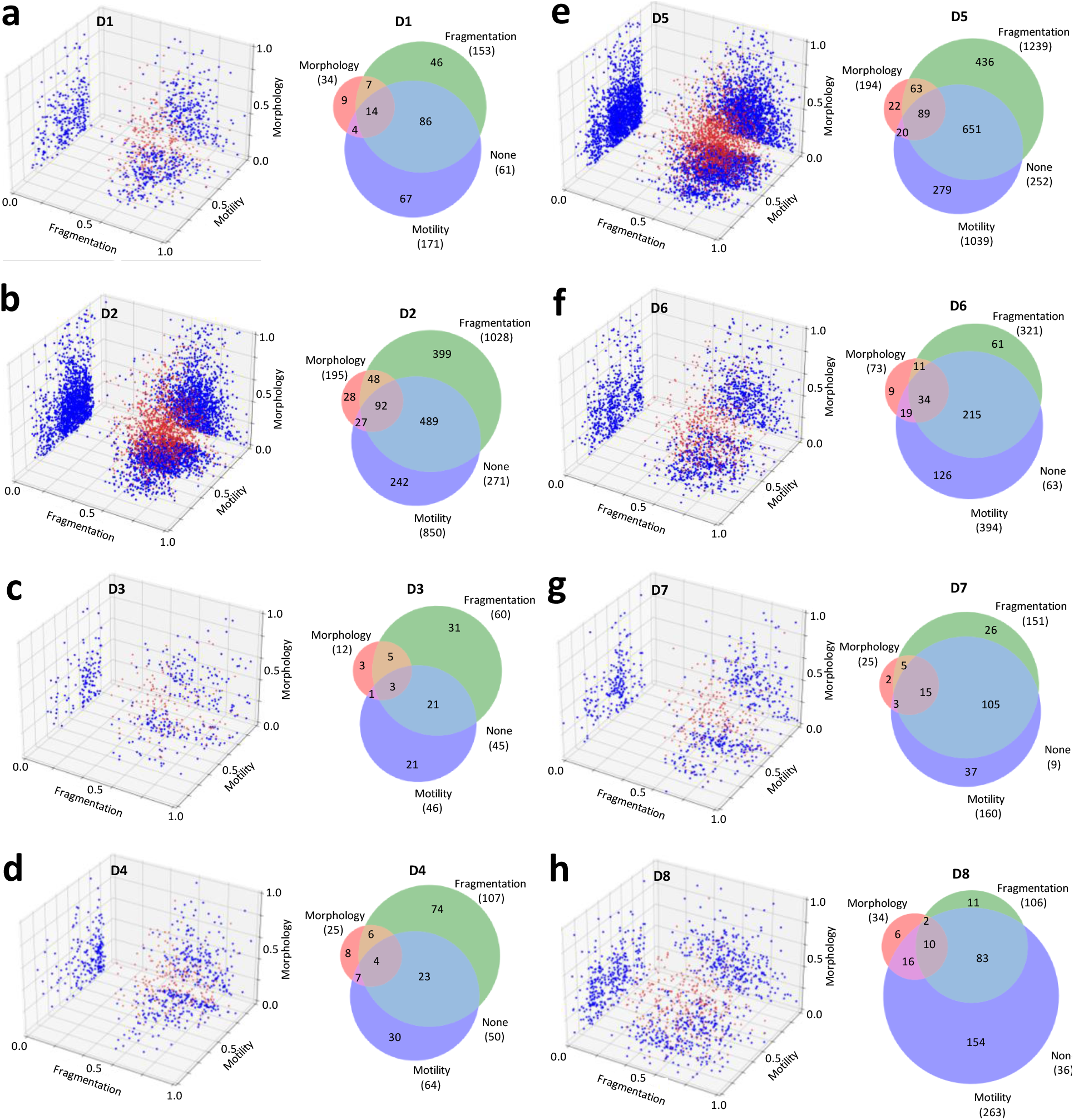
Triple-criteria sperm evaluation at the single-cell-level for eight donors. (Left) 3-D scatter plot, where each cell is represented by a red dot, and its three projections onto the 2-D planes are represented by blue dots. (Right) Venn diagrams for the donors with each of the three fertility criteria and their intersections. (morphology above a threshold of 0.5, motility above a threshold of 0.29, and fragmentation above a threshold of 0.61). a) Donor 1. b) Donor 2. c) Donor 3. d) Donor 4. e) Donor 5. f) Donor 6. g) Donor 7. h) Donor 8.

To check whether the tests dependencies were consistent between the donors, we checked the statistical independence between the criteria per each donor. This was implemented by multiplying the three percentages of cells passing each criterion and comparing the result to the percentage of the cells passing all three criteria. For example, for Donor 3 (**Figure 5c**), the number of cells that actually pass all three criteria is 1.53 times of the number of cells that are expected under the independent-criteria assumption, whereas for Donor 4 (**Figure 5d**), the same ratio is only 0.95. This shows that the criteria are not completely independent, as the expected percentage of passing cells under the independent-criteria assumption is different from the end-result and that the criteria dependencies differ between the donors.

We now define the fertility score based on the current practice and two new fertility scores based on the presented triple-criteria capability. Due to the inability to measure morphology, motility, and DNA fragmentation all together on the same cell, fertility scoring today is carried out by checking each criterion separately, on different parts of the sample, and either assuming that the sample is homogenous or repeating the examination to check for consistency. The current fertility assessment relies on averaging the cells tested per each criterion separately, resulting in three different scores, yet missing the intersections between the criteria. To define the current-practice score, we first calculated the average results of each fertility assessment (morphology, motility, and DNA fragmentation) separately, averaging over all cells per donor. Then, we normalized each assessment score over all donors and found the distance of the resulting (*x, y, z*) point from the origin of the 3-D space. We define *P* as the distance from the origin in the 3-D space, where each dimension corresponds to one of the three normalized coordinates (morphology (*x*), motility (*y*), and DNA fragmentation (*z*)):

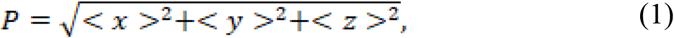

where 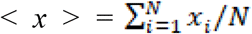 is the normalized morphology value averaged for all *N* cells measured per donor, 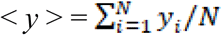 is the normalized motility value averaged for all *N* cells measured per donor, and 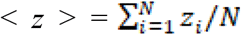 is the normalized DNA fragmentation value averaged for all *N* measured cells per donor. This normalized overall score *P* represents the fertility scores obtained by the current practice, as the averaging was done on each criterion separately, with the underlying assumption of sample homogeneity.

On the other hand, based on the presented unique capability to measure triple criteria per cell, we can define a new score, *K*, as follows:

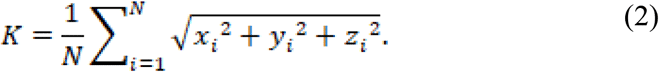

This parameter takes the distance from the origin for each cell in the normalized 3-D space, and averages all these distances per donor.

We also define *K*_1_ as the percentage of cells that pass a certain threshold set of all three criteria, per donor, where is the logical ‘and’ operator. For example, in the case of **Figure 5**, it is defined as follows:

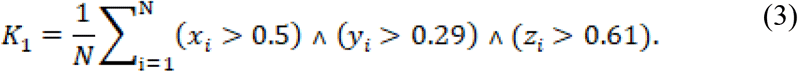

These new scores, *K* and *K*_1,_ depend on how well the cells stand in all three criteria, whereas a donor might have no sperm cells that pass all three criteria together and still have a high previous fertility score *P*.

We now show that these new scores result in different donor grading, implying that erroneous donor grading is done today, even under the homogenous sample assumption. **Figure 6** shows the previous fertility scores together with the new scores obtained for the eight donors. In this figure, each sphere represents a donor. The distance of the center of the sphere from the origin of the 3-D space represents the donor’s status as per the current practice, *P*, and can be seen as the first value next donor number. On the other hand, the sphere diameter is correlative to the donor’s first new fertility score, *K*, which can be seen as on the right in the second row next to each donor’s sphere. The second new score, *K*_1_, can be seen on the left in the second row next to each donor’s sphere. As seen, the donors’ rankings according to the old and new scores differ from each other. Donor 3, for instance, takes the last place according to the *K* score, as this donor’s average sperm quality is the lowest, although Donor 4 would be labelled as the worst donor according to the previous fertility grading, *P*. Donor 1 takes the fourth place according to the *K* score, while he takes fifth place according to the *K*_1_ score and sixth place according to current score, *P*. On the other hand, the two best donors, Donors 7 and 6, are consistent among all three scores, while Donor 5 is the third best donor for both the *K* and the *P* scores but not for the *K*_1_ score, implying that the new scores may be especially useful in discriminating situations of infertility.

**Figure 6.**
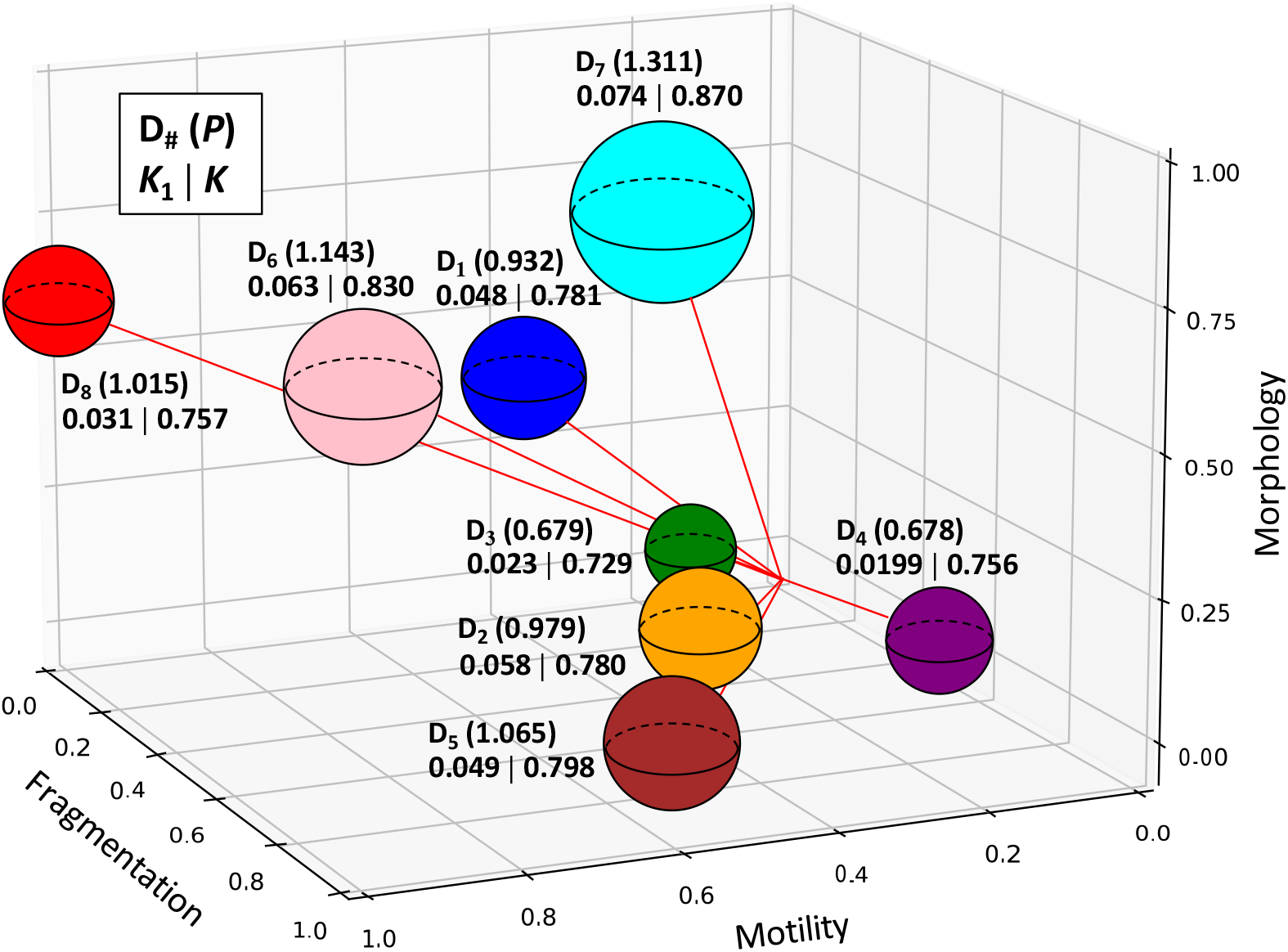
Donor fertility scoring. 3-D plot representing all donors by their fertility score based on the current-practice score (in brackets, after the donor number) and the proposed scores *K*_1_ | *K*, based on having the ability to measure all criteria together at the individual-cell level. The distance of each donor from the origin is correlative to the current-practice fertility score, whereas the size of each sphere is correlative to the proposed score.

## 3. Conclusion

In this work, we present a new capability to simultaneously measure individual sperm morphology, motility and fragmentation, as well as virtually stain the sperm cells. Today, a clinician is unable to gather this necessary information per sperm cell, and instead the fertility examination provides independent percentages regarding each evaluation separately, on different parts of the cell population, where statistical changes may occur between sub-samples. Thus, currently, there is no information regarding how many cells would have passed two or all three tests. In addition, the harsh fixation and staining protocols required for performing the morphology and DNA fragmentation assays might damage the reliability of the assays if not performed correctly, and create different results when performed by different labs. Our virtual classifiers solve these issues as they do not require staining and perform all three assays on the same cells, while they are alive and dynamic. We show that our virtual assays correlate with the chemical gold standards, and when applied together on each cell, give us the capability to better analyze the fertility status of a patient without missing the intersections between criteria. We show that patients may differ in the statistical dependencies of the different tests (**Figure 5**) and that consequently a patient with overall lower independent scores may very well have a higher score per cell or even a higher percentage of qualified sperm cells (**Figure 6**), emphasizing the need for the new approach and the insufficiency of the current examination.

This approach is expected to give rise to personalized sperm quality evaluations by adjusting the thresholds to suit the current need. **Figure S5** and **Video S2** show how different thresholds result in different sperm groups. Without any specialized needs, one may want to use the original thresholds, and obtain a result similar to that which is shown in **Figure S5a**. However, another would possibly want to increase the fragmentation threshold, as is shown in **Figure S5b**, securing a low percent of fragmented cells from a potential sperm donation. In another case, one may personalize these thresholds for selecting sperm cells for fertilization and may want to raise all thresholds in order to select the most promising sperm cells, creating scatter plots similar to the one in **Figure S5c**. Other situations may result in decreasing or increasing different thresholds, depending on which patient is being evaluated, what types of infertility problems he possesses and for what reason this evaluation is being held.

Here, we demonstrated the usefulness of the method for fertility evaluation prior to IVF. In the future, the integration of a portable clinic-ready interferometric module that can be attached to the standard microscopic systems used today in the clinic, together with our virtual staining and classification triple-criteria method at the single-cell level, can enable computer-assisted sperm selection during IVF in real-time, allowing clinicians to choose the most qualified sperm cells for egg injection, with the best morphology, motility, and DNA fragmentation scores. Since the scoring is automatic, rather than manual, in the future, the system may be integrated with a robotic selector and independently choose the sperm cells with the highest probability of fertilization. Moreover, incorporating these systems will give rise to advanced research linking types of chosen sperm cells with fertilization and pregnancy success, with the ability to personalize medicine for patients suffering from fertility problems.

To conclude, we have suggested a new approach that utilizes stain-free quantitative phase imaging and deep learning, in order to improve male fertility assessments through combined morphology, motility and DNA fragmentation scores, allowing clinicians and researchers to obtain previously unattainable fertility evaluations at the single sperm-cell level.

## 4. Methods

### Sample Preparation

Upon obtaining the institutional review board approval of Tel Aviv University, semen samples were collected from eight randomly selected donors between the ages of 18 and 45, after they signed an informed written consent. The samples were left to liquefy for 30 minutes. Sperm cells were isolated from the semen fluid using a PureCeption bilayer kit (ORIGIO, Målov, Denmark) in accordance with the manufacturer’s instructions. Then, the sample was suspended in human tubal fluid (HTF, 3 mL) medium (Irvine Scientific, CA, USA) and placed on top of a 40% and 80% silicon bead gradient and centrifuged for 20 min at 1750 rpm. The supernatant was removed and cells in the pellet were resuspended in 5 ml of HTF before centrifugation at 1750 rpm for 5 min. The supernatant was discarded and the pellet containing the living sperm cells was resuspended in HTF (100 µL). For the stain-based morphological assays, 5 µL of the sample were mixed with 5 µL of QuickStain (Biological Industries, Beit Haemek, Israel). For the stain-based DNA fragmentation assays, the sample was washed with HTF (5 ml) before centrifugation at 1750 rpm for 5 minutes. The supernatant was discarded, and the pellet was vortexed gently. The cells were then fixed by slowly adding 5 ml of methanol-acetic acid mix (3:1). After 5 minutes of incubation at room temperature, the cells were centrifuged at 1750 rpm for another 5 minutes and the supernatant was discarded. The pellet was resuspended in the fixative solution (0.1 ml). Next, the solution was placed on the cover slip and stained with acridine orange (0.19 mg ml^-1^) for several hours before imaging.^[31]^

### State of the Art Fertility Examination

For each donor, 10 µL of the sample were used for stain-based morphological evaluation, 10 µL for concentration and motility evaluation, 10 µL for stain-based DNA fragmentation evaluation, and the rest was used for stain-free interferometric assays. All state-of-the-art examinations were carried out according to the WHO2010 guidelines.^[3]^ For stain-based morphological analysis, 200 QuickStain-labeled cells were assessed on a slide according to WHO2010 guidelines using an inverted microscope (CKX53, Olympus) with 100× oil objective and a 10× ocular. Sperm concentration and motility were examined in a Makler counting chamber (Sefi Medical Instruments, Haifa, Israel) under an inverted microscope (Primovert, Zeiss) with a 20× oil objective and 10× ocular. For stain-based DNA fragmentation analysis, the acridine-orange-labeled cells were imaged on a slide by a confocal fluorescence microscope (Zeiss LSM 510-META) using a 25×, 1.4 numerical-aperture microscope objective. Excitation wavelengths were 477-488 nm, and emissions were at 572-668 nm and 505-550 nm.

### Interferometric Phase Imaging

The optical system architecture is shown **Figure S1**. The sample was illuminated by a HeNe laser (Thorlabs Model HRP170) with a wavelength of 632.8 nm, and was imaged by an inverted microscope consisting of microscope objective MO (Olympus PlanApo N, 60×/1.42 oil) and tube lens TL (focal length: 150 mm). The sample magnified image entered a clinic-ready portable interferometric module,^[24]^ where it was split by beam splitter BS. One beam was spatially filtered by lenses L1 and L2 (f = 100 mm) and a 15 um pinhole, turning it into a reference beam, and then projected towards the camera by a corner retroreflector mirror, and the other beam was projected towards the camera at a small angle using a laterally shifted corner retroreflector mirror. The combined beams were projected through a 4f system comprised of lenses L3 (f = 50 mm) and L4 (f = 40 mm), and the resulting off-axis image hologram was produced on the digital camera (iDS UI-3880CP Rev.2, 3088 × 2076 pixels, 2.4 × 2.4 μm each). 231 one-minute interferometric videos were recorded at 40 frames per second on non-overlapping field of views for the eight donors. Each off-axis hologram from each video frame was digitally Fourier transformed and one of the resulting cross-correlation terms was cropped and inverse Fourier transformed, thereby reconstructing the complex wave front of the sample. The phase argument underwent a phase unwrapping,^[32]^ and the result was divided by 2π and multiplied by the illumination wavelength, resulting in the optical path delay (OPD) or optical thickness map of the sample.

### Cell Tracking Algorithm

The resulting OPD videos were processed using a specially designed cell-tracking algorithm, resulting in a space-time array per cell. This array consisted of the location of the cell in each frame and was used for the automatic motility, stain-free fragmentation, and stain-free morphological assessments. Connected component labeling is used to find the centroid of each cell. Detected cells in different frames that were close in time and space were grouped as the same cell; Cells were included if recognized in less than 5 consecutive frames, cells containing less than 15 frames were not included. The resulting dataset was comprised of 683,604 frames and 12,126 cells before further filtering.

### Automatic Stain-Free Morphological Evaluation

For morphological evaluation, the OPD maps were filtered so that only frames that resembled in-focus cells viewed from an axis perpendicular to the major axis of the cell remained. Cells with flat head orientation were chosen. We then extracted the total head area, nucleus area, acrosome area, mean posterior-anterior difference, mean OPD and variance of the OPD. The embryologist classified each cell according to the five following qualifications and provided a pass-or-fail result:^[3,33]^ (a) The head should be smooth with an oval-like shape with a width-to-length ratio of roughly 3:5. (b) The acrosome should be between 40%-70% of the head area and should be clearly visible. (c) There should be no more than two small vacuoles, and they should occupy less than 20% of the sperm head and exist outside the post-acrosomal region. (d) The midpiece should be aligned with the head’s major axis, should be about the length of the sperm head and should be regularly shaped and slender. (e) Residual cytoplasm should not exceed one third of the sperm’s head size. We used a previous sperm database^[30]^ and built here a new CNN classifier. The input to the model was the preprocessed OPD image of the cell and the output was a binary vector predicting if the cell passed each of the five criteria (a-e), and a direct overall prediction of whether the cell passed all five criteria together according to the embryologist. The architecture of the model is shown in **Figure 2**. It consisted of 3 CNN layers with 32, 64 and 128 kernels of size 3×3, ReLU activation functions and padding set to ‘same’, with Max Pooling layers of size 2×2, dropout layers with probability of 0.5 and 2 fully-connected (FC) layers of size 1500 and five with the same dropouts and activations, with an L2 kernel regularizer consisting of a penalty of 0.001 and a final Sigmoid activation function. The Adam optimizer was used with a learning rate of 0.001 and the compilation consisted of a custom binary cross entropy weighted loss. The model was trained with a batch size of 100 and 6000 epochs with a training-validation-test split of 80%-10%-10%. The code was written in Python in Google Collaboratory using Tensorflow 2.3.0 with Tensorflow Keras 2.4.0.

### Automatic Motility Evaluation

To automatically assess sperm motility, we created an algorithm that takes the space-time arrays of the cell tracking algorithm as the inputs, and outputs each donor’s motility scores. A progressive sperm cell swims fast and in a linear projection at a velocity of at least 25 μm sec^-1 [3]^ As linearity is expected only for short time-periods and short distances, one second windows in increments of half a second were selected to assess progressivity^[33]^ and the median values were chosen for each motility score per cell. The motility features extracted were: (a) Curvilinear velocity (VCL) – time averaged velocity of a sperm head along its actual curvilinear path. (b) Straight line velocity (VSL) – time averaged velocity of a sperm head along its linear estimated path – a straight line connecting its first and last spatial points. (c) Average path velocity (VAP) – time averaged velocity of a sperm head along its average estimated path. The average path was calculated as a third-degree polynomial approximation of the original path. (d) Linearity (LIN) – the linearity of the curvilinear path calculated as the ratio of the VSL to the VCL. (e) Wobble (WOB) – oscillation of the actual path about the average path, calculated as the ratio of the VAP to the VCL. (f) Straightness (STR) – linearity of the average path, calculated as the ratio of the VSL to the VAP. (g) Beat-cross frequency (BCF) – the average rate at which the curvilinear path crosses the average path. (h) Mean angular displacement (MAD) – the time-averaged absolute values of the instantaneous turning angle of the sperm head along its curvilinear trajectory. (i) Progressiveness (PROG) – A cell is considered progressive if its VSL is at least 25 μm sec^-1^ and its LIN is at least 0.6. To compare our results to the previous qualitative motility results according to the WHO, we created a least-squares-based linear transformation that uses all nine features (a-i) and maps them to a prediction in the original scale of 1-3 (immotile to progressively motile) that were given to the cell by the embryologist. After all cells were given a motility score based on this three-class assessment, the average was taken as the final motility score per donor.

### 4.7 Stain-Free DNA Fragmentation Evaluation

We created neural networks for DNA fragmentation classification and virtual staining. For training, we used the database described in our previous work, which contains pairs of OPD maps and the same cell labeled in acridine orange and imaged under a confocal fluorescence microscope (Zeiss LSM 510-META).^[31]^ Each fluorescent cell was evaluated in a random order by the experimentalist in a color scale of 1 (red) to 5 (green). This number was used as the gold standard while training a five-class CNN classifier. The inputs to the network were the stain-free OPD map, and the cell dry mass, mean anterior-posterior difference, head, acrosome and nucleus areas and head OPD mean and variance. The CNN classifier architecture is shown in **Figure 4a**. The OPD image went through five CNN layers while the features went through three FC layers. All the layers in the first two parts had the ReLU activation function. The result of both the sub-networks were concatenated and inserted into the third and final part consisting of an additional three FC layers and the output, while the activation function of the hidden layers was the Leaky ReLU with a slope coefficient (of 0.3, and the final activation was a sigmoid with the loss being weighted binary cross-entropy. The ground truth of each cell was between zero and one, while class 1 was given a value of 0.2, class 2 – a value of 0.4 and so on with class 5 being given a value of 1. The predicted class of each cell was the one whose value was closest to the model prediction. The classifier had a train-validation-test data split of 80%-10%-10%. The model was trained on five different data folds with augmentation and class weights. Another network was used for individual cell virtual staining. It mapped the predicted fragmentation status from the classifier to the expected acridine orange color and visualized the cells as though they have been stained by acridine orange, but without using chemical staining. First, we took all the fluorescent images of each cell and split the cell bounding box into nine three by three patches. For each patch, we calculated the ratio between the average green and red channel values. Then, for each cell, we created a nine-feature vector of the results sorted in an ascending order. We used a k-means algorithm (k = 9) to cluster the images, and then grouped the clusters into five groups of colors. Then, we created a GAN, with architecture shown in **Figure 4b**, which was trained separately for each class, as the input to the discriminator would switch from a resized version of the original image belonging to that class, to the generated image.

## Author Contributions

KBY developed the main ideas, designed the algorithms and experiments, implemented the algorithms, conducted the experiments, processed the data, and wrote the paper. SKM built the optical system, designed part of the image-processing algorithm, conducted the experiments, and wrote part of the paper. ML developed the main ideas, prepared the biological samples, conducted the experiments, and wrote part of the paper. IB prepared the biological samples, conducted the experiments and wrote part of the paper. SK and IM implemented the morphology classifier algorithms. DB and RCC implemented the motility algorithms. YNN designed the morphology virtual-staining algorithm. NTS conceived the project, developed the main ideas, designed the algorithms and experiments, wrote the paper, and supervised the project.

## Supporting Information

**Figure S1.**
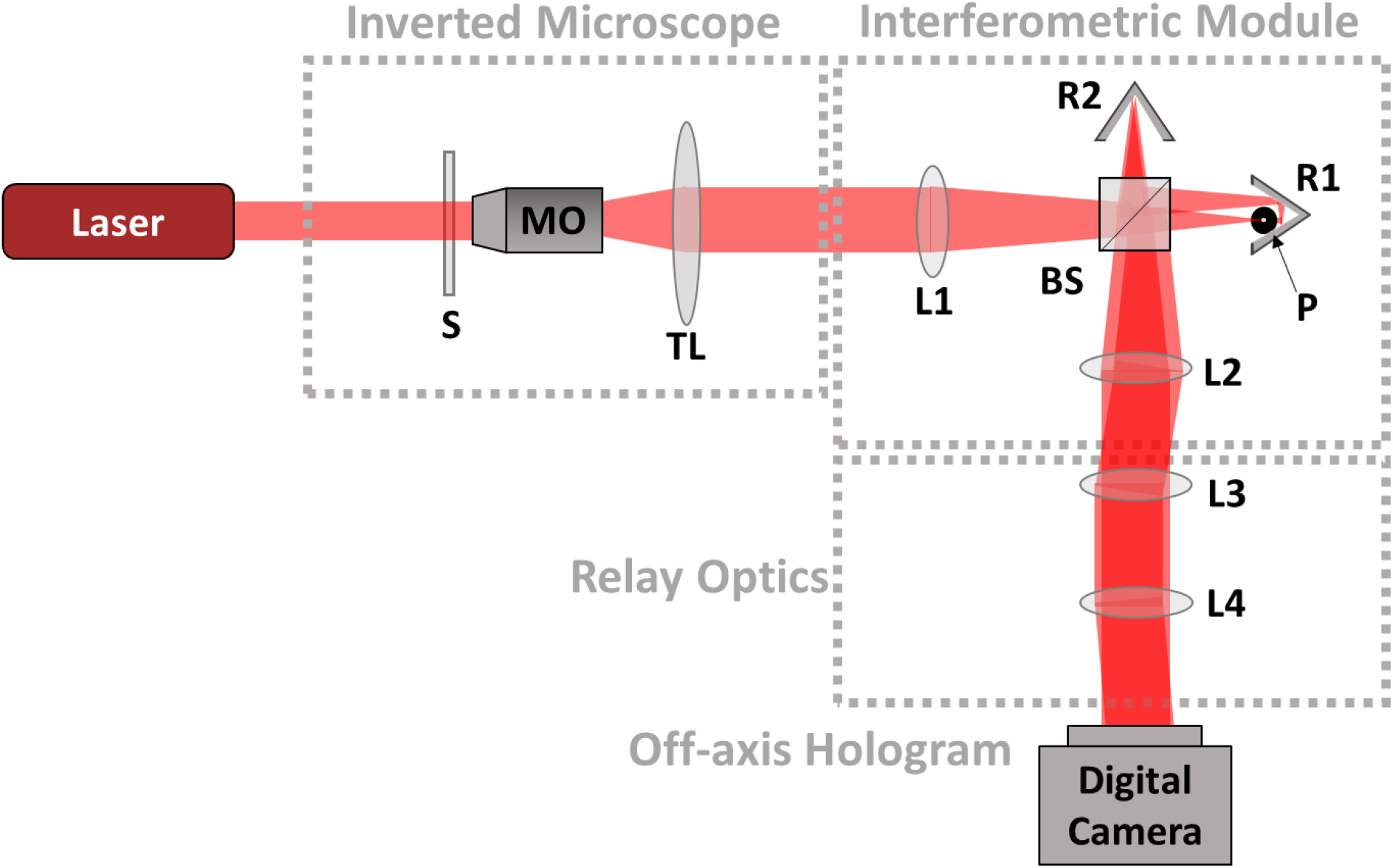
Diagram of internal design of the optical system. S, sample; MO, microscope objective; TL, tube lens; L1, L2, L3, L4, lenses; BS, beam splitter; RR1, RR2, retro-reflector mirrors; P, Pinhole.

**Figure S2.**
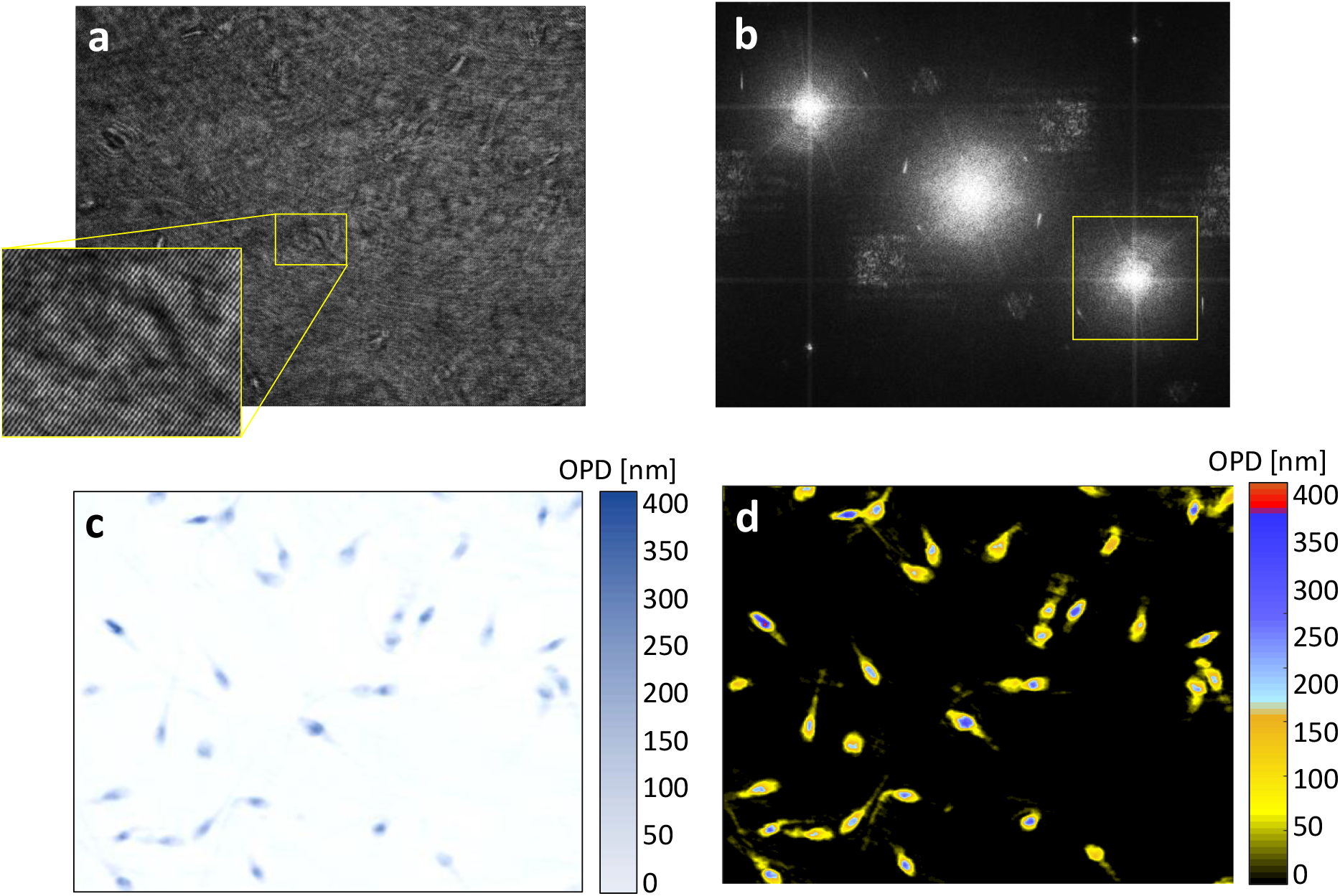
Dynamic quantitative phase imaging of rapidly swimming sperm cells. a) Off-axis hologram acquired by the digital camera without chemical staining in the full frame rate of the camera. b) Spatial power spectrum, with the selected cross-correlation term marked by a yellow square. c) The stain-free OPD (optical thickness) profile obtained by processing the cropped cross-correlation term. d) Parametric coloring of the OPD profile, resembling HEMA chemical staining. See dynamics in **Video S1**.

**Figure S3.**
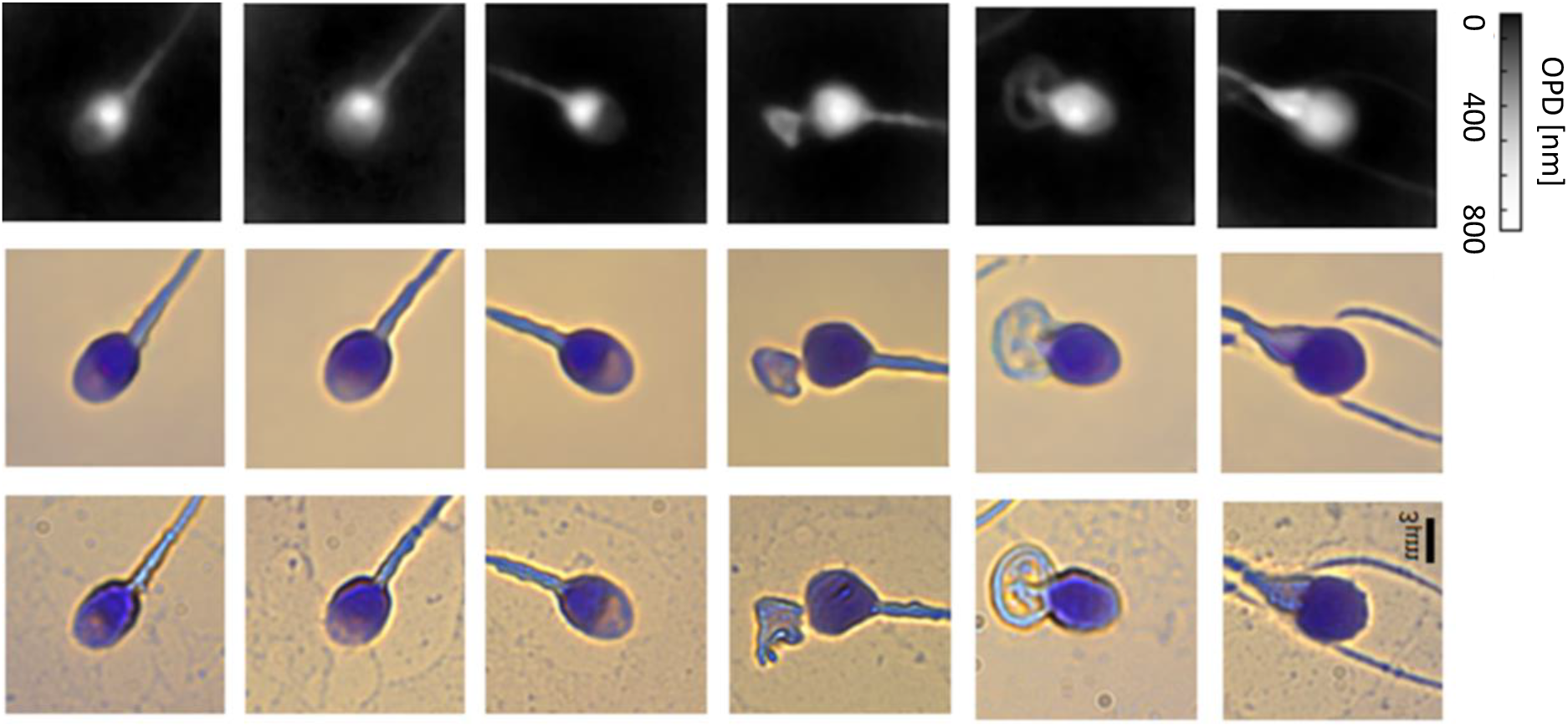
Morphological virtual staining of individual sperm cells by deep learning. We used the stain-free quantitative phase profiles of the cells as inputs to a neural network that performed the virtual-staining mapping, after being trained with the coinciding chemical staining images. The first row shows the quantitative phase images extracted from the single holograms. The second row shows the coinciding virtual stained images, generated by the network. The third row shows the coinciding bright-field chemically stained images of the same sperm cells that the network did not see; yet it could present the cells as if they were stained, as shown in the second row, based only on the stain-free quantitative phase images shown in the first row. The first three columns show normal morphology cells, and the last three columns show pathological cells.

**Figure S4.**
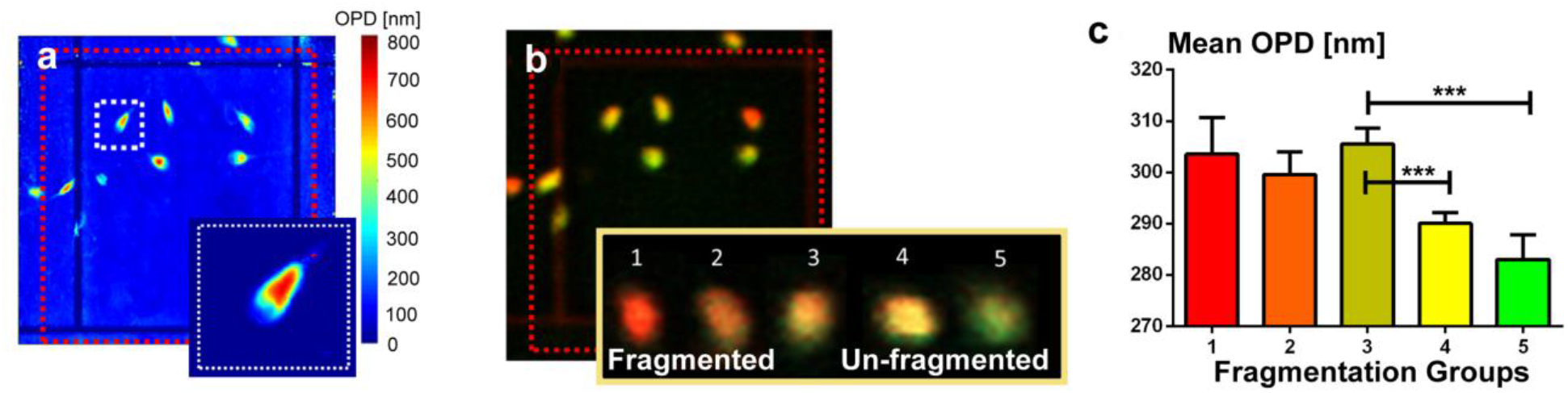
Stain-free DNA fragmentation measurements, used as the database of the deep learning classifier. a) Sperm OPD map of sperm cells obtained without staining. b) The same sperm cells, later chemically stained by acridine orange, indicating the sperm cells with fragmented DNA by the emitted color. c) Mean OPD of different DNA fragmentation groups (*** p value < 0.001). A database of pairs of images, example of which are shown in **a** and **b**, have been used for training the deep neural networks that can classify live cells by their fragmentation group without chemical staining and for DNA-fragmentation virtual staining.

**Figure S5.**
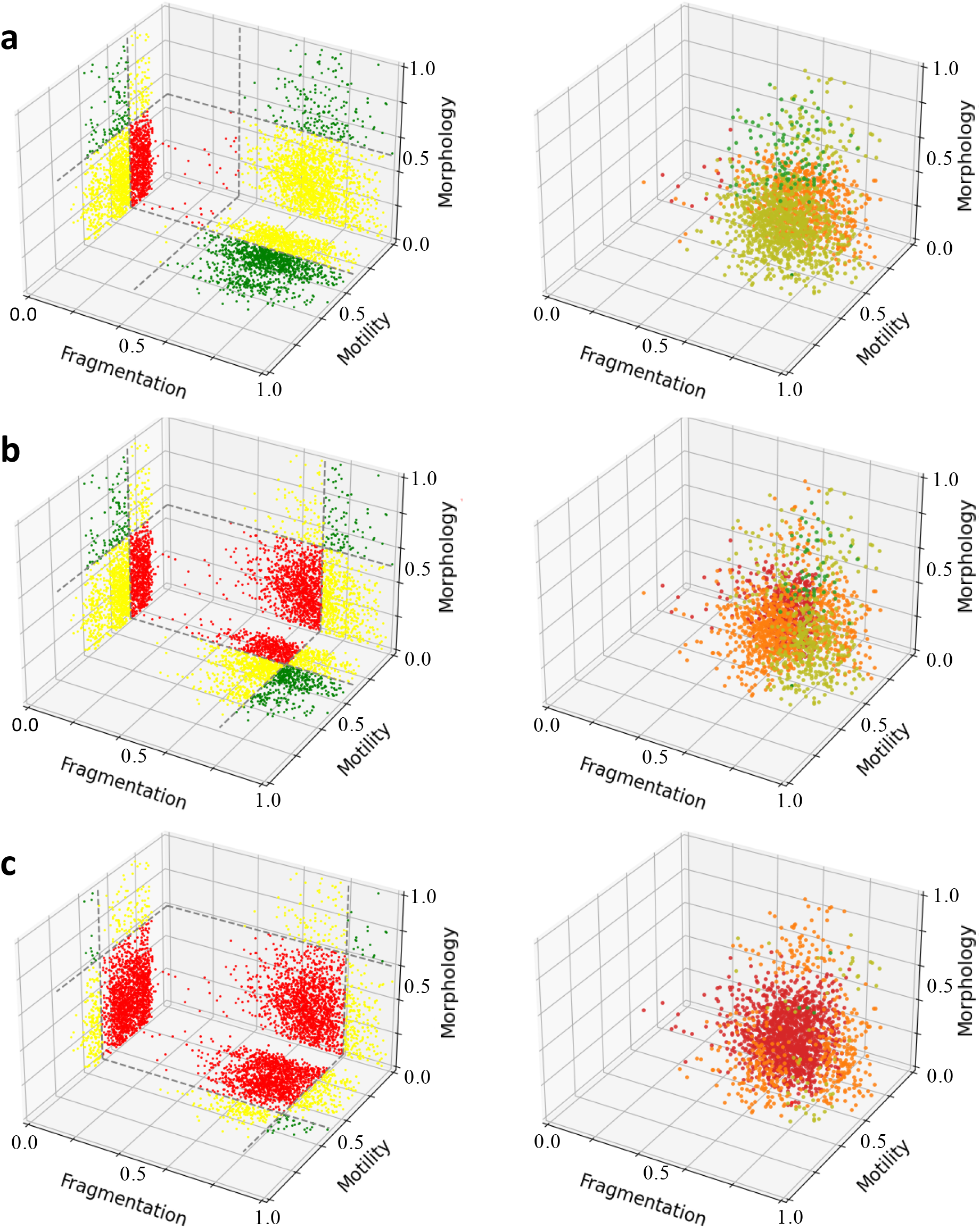
Cell classification for Donor 5 using different thresholds. Left: Projections on the 2-D planes. Green cells pass both criteria on the plane, yellow cells pass only one criterion, and red cells pass none. Right: 3-D locations of each cell. Green cells pass all criteria and are qualified as good sperm cells, olive cells pass only two out of the three criteria, orange cells pass only one and red cells fail in all. a) Morphology, motility and fragmentation thresholds of 0.5, 0.29 and 0.3, respectively. b) Thresholds of 0.5, 0.29, 0.7. respectively. c) Thresholds of 0.6, 0.5, 0.8, respectively. See dynamic threshold changing in **Video S2**.

